# Extinction drives a discontinuous temporal pattern of species-area relationships in a microbial microcosm system

**DOI:** 10.1101/2022.11.11.516089

**Authors:** Wei Deng, Na Li, Chao-Zhi Zhang, Rui An, Xiao-Yan Yang, Wen Xiao

## Abstract

The species area relationship (SAR) can be a general law in ecology only if its formation mechanisms are clarified. Essentially, the SAR addresses the relationship between regional area and biodiversity, which is shaped by processes of speciation, extinction and dispersal. Since those three processes has temporal dynamics, we propose the hypothesis that the occurrence of SAR should also have temporal dynamics. Here, we designed independent closed microcosm systems, in which dispersal/speciation can be excluded/neglected, to reveal the role of extinction in shaping the temporal dynamics pattern of SAR. We find that extinction can shape SAR in this system independently. Due to the temporal dynamics of the extinction, SAR was temporally discontinuous. We also observe that small-scale extinctions modified community structure to promote ecosystem stability and shaped SAR, while mass extinction pushed the microcosm system into the next successional stage and dismissed SAR. Our result suggested that SAR could serve as an indicator of ecosystem stability; moreover, temporal discontinuity can be the explaination for many controversies in SAR studies.

## Introduction

The species-area relationship (SAR) is the most studied and well-documented pattern of biodiversity and is considered one of the few “laws” in ecology ^1–4^. The SAR describes the positive correlation between species richness and habitat area, which has been widely used in biodiversity prediction and species extinction assessment and has played a fundamental role in understanding biodiversity patterns, species conservation planning and even the construction of biodiversity theory ^5–8^. There are currently three dominant hypotheses regarding the formation of SARs: 1) decrease in area leads to an increase in extinction (the area per se hypothesis) ^9–12^, 2) the increase in habitat heterogeneity due to the increase in area (the habitat diversity hypothesis) ^13,14^, and 3) the increase in the number of individuals observed as the area increases (the passive sampling hypothesis) ^15^. In addition, edge effects and resource concentration are also thought to have played a key role in the formation of SARs ^16^. Of course, some researchers believe that the SAR results from a combination of the above effects ^17^. However, to date, the relative contributions and exact operation mode of many SAR hypotheses have not been determined ^18^. Thus, the mechanism underlying SAR formation is still unclear. Clarifying this mechanism is necessary for the SAR to truly become an ecological law. Connor & McCoy (1979) argue that determining the mechanism of the SAR requires direct validation of a hypothesis through experimental design and exclusion of other factors ^17^. For example, to conclude that habitat diversity is the only mechanism underlying an SAR, it is necessary not only to directly prove the effect of habitat diversity on species richness but also to prove that there is no relationship between extinction rate and area.

Essentially, the SAR addresses the relationship between regional area and biodiversity, which is shaped by processes of species generation, extinction and migration. With the exception of the passive sampling hypothesis, each SAR hypothesis posits that the SAR is shaped by extinction; the widely accepted area per se hypothesis in particular emphasizes the effect of extinction. Therefore, to clarify the formation mechanism of the SAR, extinction should be a research focus. To determine the independent mechanism of extinction, an experimental design that excludes many influencing factors, such as migration, habitat heterogeneity, resource availability, biodiversity formation background ^19^, and sampling effects, is needed.

In this study, an independently sealed microcosm system was constructed with homogeneously mixed Chinese pao cai (fermented vegetables) soup and opaque closed culture flasks. The use of amplicon sequencing methods to conduct a comprehensive assessment of microbial diversity in the microcosm circumvents the impact of sampling effort. In this system, the historical background of microbial diversity is consistent, the environment is homogeneous, resource density is consistent, and the impacts of migration are completely avoided. The process of species generation in this system can also be ignored, meaning that extinction should be the dominant force shaping the biodiversity of the microcosm system. Studies have found that extinction is a dynamic process. Therefore, we continuously monitored the biodiversity of microcosm systems at 26 time points over a 60-day period. We hypothesised the following: 1) In the microcosm systems, extinction would be the main mechanism the SAR; 2) extinction in the microcosm would be temporally dynamic, and phased extinction would promote the succession of microcosmic ecosystems; and 3) consistent with the successional phase of the community, there would also be temporal dynamics in the occurrence of SARs.

## Materials and Methods

### Preparation of the pao cai soup

First, 35 kg of white radish (*Raphanus sativus*), 35 kg of cabbage (*Brassica oleracea*), 2 kg of chili pepper (*Capsicum frutescens*), 1 kg of ginger (*Zingiber officinale*), 1 kg of peppercorns (*Zanthoxylum bungeanum*), 2.5 kg of rock sugar, and 210 kg of cold boiled water (containing 6% salt) were divided into six ceramic jars. After 7 days of natural fermentation at room temperature, the pao cai was filtered out with sterile gauze to obtain 200 kg of pao cai soup. To ensure an even distribution of microorganisms in the soup, the soup was mixed well and then left to rest for 12 h, the supernatant was taken, and the soup was left to rest for 12 h again.

The plants used in this study were cultivated vegetables which purchased from the vegetable market at the study site. All local, national or international guidelines and legislation were adhered to in the production of this study.

### Establishment of the microcosm system

Seventy-eight 10 ml, 20 ml, 50 ml, 100 ml, 250 ml, 500 ml, and 1000 ml sterile glass culture flasks were filled with pao cai soup, the bottle mouth was sealed with sterile sealing film, and the bottle was capped without leaving any air (Fig. S1). Each flask became a microcosm and was cultured in a 25 °C incubator.

### Sample collection

Before the microcosm system was established, a sample of well-mixed pao cai soup was taken as a reference to establish background biodiversity. Samples were collected daily for 1-10 day after the establishment of the microcosm and then collected every 2 days for 10-30 day and every 5 days for 30-60 day. Three different microcosms of the same volume were established. Monitoring was carried out for 60 days, and a total of 546 samples of 7 volumetric gradients were obtained at 26 time points. At the time of sampling, the pao cai soup in the microcosm was mixed, and 50 ml of sample (10 ml of sample was collected for microcosm systems with a volume of less than 50 ml) was collected. The sample was centrifuged at 8000 rpm for 10 min, the supernatant was collected for pH determination, and the pellet was stored in a −80 °C freezer.

### Microbial analyses

Microbial DNA was extracted from pao cai samples using the E.Z.N.A.® Soil DNA Kit (Omega Biotek, Norcross, GA, U.S.) according to the manufacturer’s protocols. For bacteria, we targeted the V3-V4 region of the 16S ribosomal RNA (rRNA) gene, using the 338F (5’ - ACTCCTACGGGAGGCAGCAG - 3’) and 806R (5’ - GGACTACHVGGGTWTCTAAT - 3’) primer pairs. For fungi, we targeted the ITS1-1F region of the nuclear ribosomal internal transcribed spacer region (ITS rDNA) gene, using ITS1-1F-F (5’ - CTTGGTCATTTAGAGGAAGTAA - 3’) and ITS-1F-R (5’ - GCTGCGTTCTTCATCGATGC - 3’). PCRs were performed in triplicate in a 20 μL mixture containing 4 μL of 5× FastPfu Buffer, 2 μL of 2.5 mM dNTPs, 0.8 μL of each primer (5 μM), 0.4 μL of FastPfu Polymerase and 10 ng of template DNA. The PCR program for the 16S rRNA gene was as follows: 3 min of denaturation at 95 °C; 27 cycles of 30 s at 95 °C, 30 s of annealing at 55 °C, and 45 s of elongation at 72 °C; and a final extension at 72 °C for 10 min. For the ITS1-1F region, the PCR program was as follows: samples were initially denatured at 98 °C for 1 min, followed by 30 cycles of denaturation at 98 °C for 10 s, primer annealing at 50 °C for 30 s, and extension at 72 °C for 30 s. A final extension step of 5 min at 72 °C was added to ensure complete amplification of the target region. The resulting PCR products were extracted from a 2% agarose gel, further purified using the AxyPrep DNA Gel Extraction Kit (Axygen Biosciences, Union City, CA, USA) and quantified using QuantiFluor™-ST (Promega, USA) according to the manufacturer’s protocol.

Purified amplicons were pooled in equimolar amounts and paired-end sequenced (2 × 300) on an Illumina MiSeq platform (Illumina, San Diego, USA) according to standard protocols. The analysis was conducted by following the “Atacama soil microbiome tutorial” of QIIME2 docs along with customized program scripts (https://docs.qiime2.org/2019.1/). Briefly, raw data FASTQ files were imported in an appropriate format for the QIIME2 system using the qiime tools import program. Demultiplexed sequences from each sample were quality filtered, trimmed, denoised, and merged, and then the chimeric sequences were identified and removed using the QIIME2 dada2 plugin to obtain the feature table of amplicon sequence variants (ASVs) ^20^. The QIIME2 feature-classifier plugin was then used to align ASV sequences to the pretrained GREENGENES 13_8 99% database (trimmed to the V3-V4 region bound by the 338F/806R primer pair for bacteria) and UNITE database (for fungi) to generate the taxonomy table ^21^. Any contaminating mitochondrial and chloroplast sequences were filtered using the QIIME2 feature-table plugin. Rare ASVs that accounted for less than 0.001% of the total sequences were excluded.

### Data analysis

Species richness is equal to taxon number, which is equal to the total number of all bacterial and fungal ASVs. The vegan package in R was used to calculate the species richness of each sample based on the ASV feature table ^22^. Using vial volume instead of area, SAR fitting was performed using a semi-logarithmic model, and its significance was tested. The semi-logarithmic model is the function S = c + b*logA, where S is species richness, A is area (in this case, volume is used instead), and b and c are fit parameters ^23^.

A microcosm system that could not be successfully amplified due to the extinction of microorganisms was recorded as an annihilation microcosm, and the microcosm annihilation rate at each time period was calculated. Using the samples on the day the microcosm was established as a reference, R was used to calculate the number of ASVs that went extinct in each system compared to the reference samples, and the extinction rate was calculated. Pearson correlation analysis was applied to examine the relationship between volume and extinction rate at each time point. Non-linear regression with a bell-shaped form was performed with time as an independent variable and pH and annihilation rate as dependent variables, and regression lines were plotted. According to the taxonomy table, bacterial ASVs were divided into acid-producing and non-acid-producing categories, and their extinction rates were calculated separately.

The agricola, ggplot2, vegan and ggpubr packages were used to draw alpha diversity box plots and perform the Wilcoxon rank sum test for differences between groups ^22,24–26^. Non-metric multidimensional scaling (NMDS) analysis was performed with the vegan package based on Bray–Curtis dissimilarity.

In addition, the potential Kyoto Encyclopedia of Genes and Genomes (KEGG) orthologue (KO) functional profiles of microbial communities were predicted with PICRUSt ^27^. Resistance-related genes were screened using the gene function predictions. The relationship between the relative abundance of resistance-related genes and the volume of the microcosm was analysed by Pearson correlation, and a forest map was plotted to present the results.

## Results

A total of 19,637,820 and 6,553,883 high-quality sequences were obtained for bacteria and fungi across all samples, respectively, with an average of 37,838 ± 4,849 (bacteria, mean ± SD) and 18,942 ± 17,186 (fungi, mean ± SD) sequences detected per sample. After removal of rare amplicon sequence variants (ASVs), we obtained 783 and 889 ASVs for bacteria and fungi, respectively.

### The temporal dynamics of SARs

During the 60 days of monitoring, a significant SAR occurred in the microcosm system in only 3-5 day, day 7, and 22-30 day (Fig. 1 c, d, e, g, p, q, r, s, t; Table S1). At 35 and 55 day, species richness in the microcosm system decreased significantly with increasing volume (Fig. 1 u, y). At the remaining time points, there was no SAR in the microbial community in the closed system (Fig. 1 a, b, f, h, i, j, k, l, m, n, o, v, w, x, z).

**Fig. 1.**
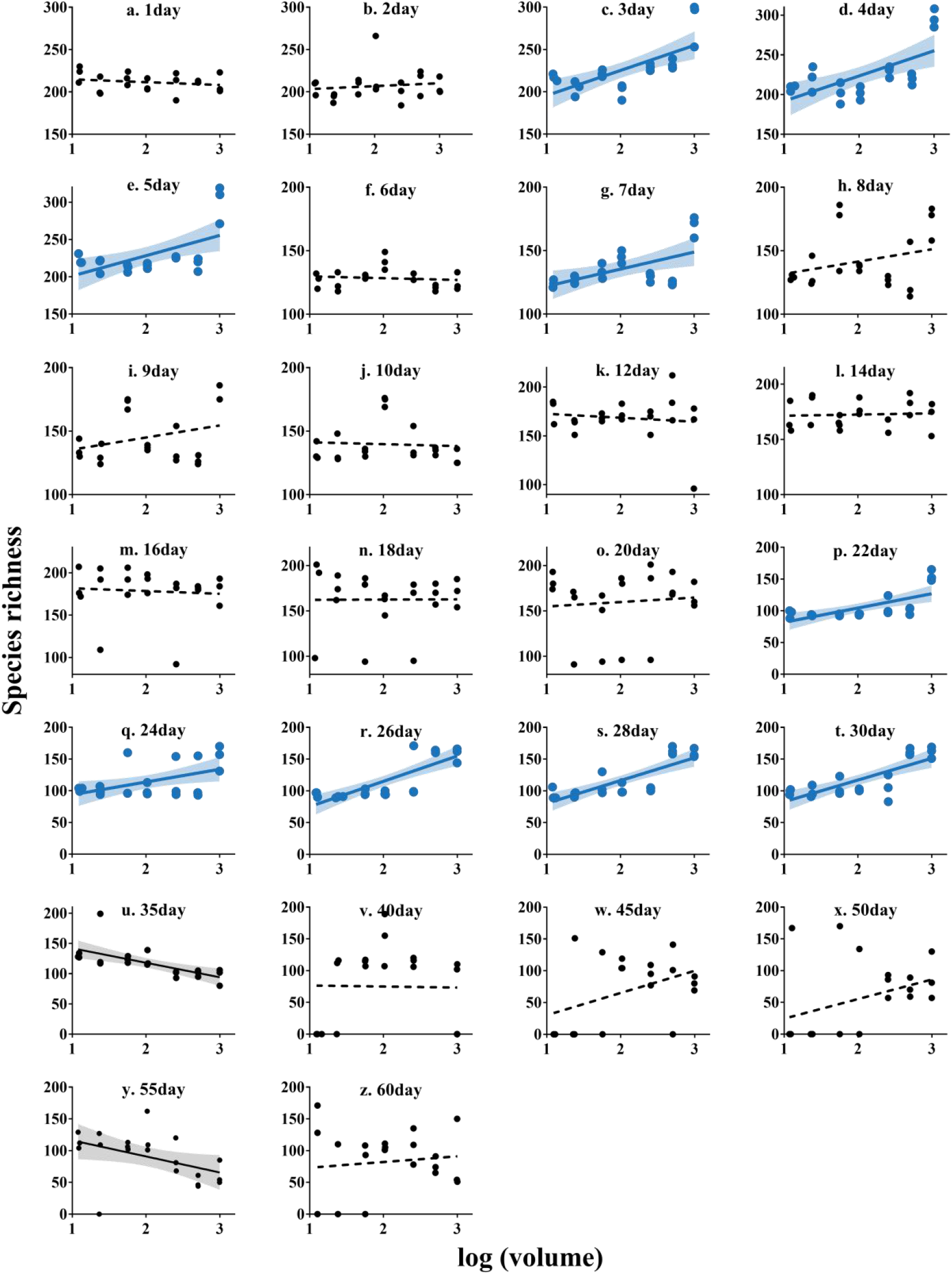
Microbial SAR at 60 days in the microcosm system. The regression line is from a linear model (volume is normally converted to the log scale). A solid line indicates a significant correlation, and a dashed line indicates that the correlation is not significant. A blue band represents the 95% confidence interval of a positive SAR, and a grey band represents the 95% confidence interval of a negative SAR (see Table S1 for linear regression results).

### Extinction drove the microbial SAR in the microcosm system

Except on day 7, the extinction rates of ASVs during the periods with an SAR were significantly inversely correlated with the volume of the microcosm (Fig. 2, Table S2).

**Fig. 2.**
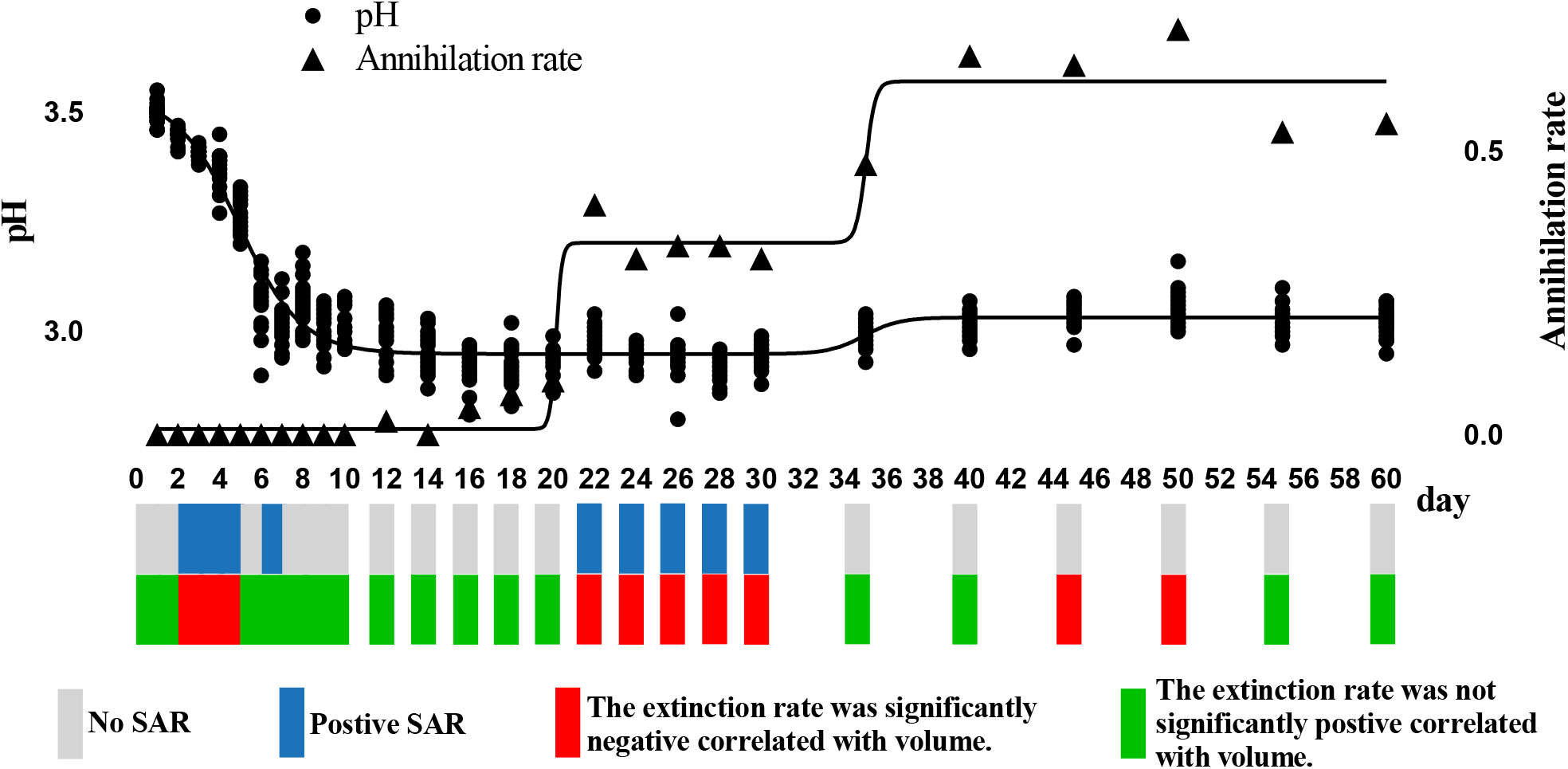
Sixty-day pH values and annihilation rates of the microcosm system, the SAR at each time point, and the relationship between extinction rate and volume. The regression line is from the model with a bell-shaped form (pH: p = 0.048, R^2^ = 0.927; annihilation rate: p = 0.045, R^2^ = 0.976). A grey block indicates that there is no microbial SAR at the corresponding time point, while a blue block indicates that there is. A green block indicates that the extinction rate is not negatively correlated with volume, while a red block indicates that the extinction rate is significantly negatively correlated with volume (see Table S2 for statistical test results).

### Phased succession of the microcosm system

The pH in the microcosm system showed a phased change. Specifically, it decreased rapidly on 1-10 day, remained stable on 12-35 day, and recovered on 40-60 day. Accordingly, the microcosm system can be divided into three phases: Phase 1 (1-10 day), Phase 2 (12-40 day), and Phase 3 (45-60 day).

In line with this finding, annihilation of the microcosm system did not occur in Phase 1, and there was systematic annihilation in Phase 2, with the annihilation rate exceeding 50% in Phase 3. The extinction rate at each volume of the microcosm system showed a temporal dynamic change pattern. The extinction rate increased significantly in the system on day 6 and day 40, and a mass extinction event occurred. The average extinction rate per unit volume in Phase 1 was 43.24% ± 13.07, the average extinction rate per unit volume in Phase 2 was 56.59% ±8.25, and the average extinction rate per unit volume in Phase 3 was 82.42% ±10.24 (Fig. 3).

**Fig. 3.**
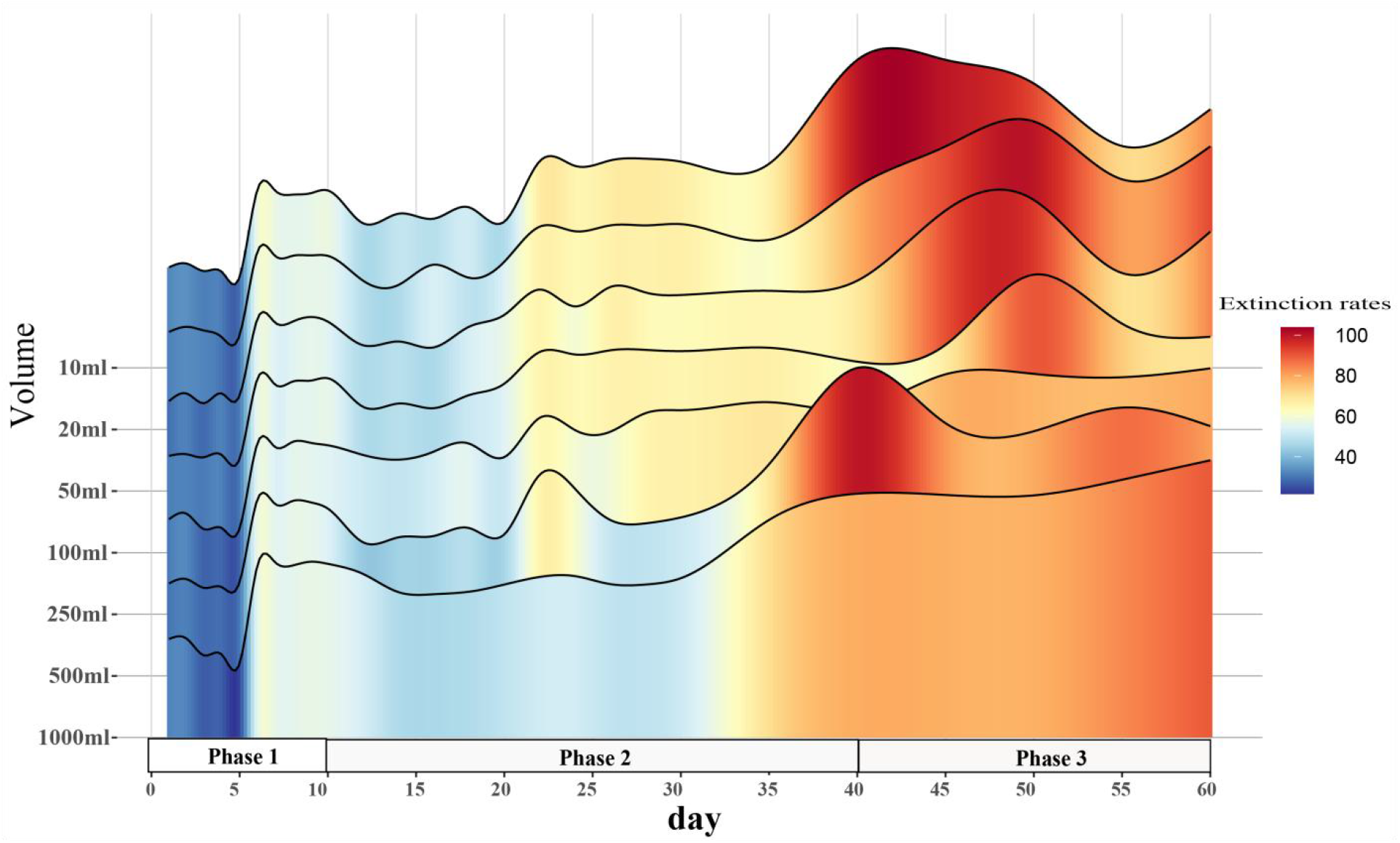
Dynamic change in the extinction rates of microcosm systems. Colours from blue to red represent a low to high extinction rate.

Corresponding to the three phases of the microcosm system, the alpha diversity and beta diversity of the microbial community also exhibited phased differentiation. The Shannon diversity index differed significantly over the three phases (p< 0.0001, Fig. 4, a). Non-metric multidimensional scaling (NMDS) analysis showed that the microbial communities exhibited phased differentiation over the three phases (stress = 0.087, Fig. 4, b).

**Fig. 4.**
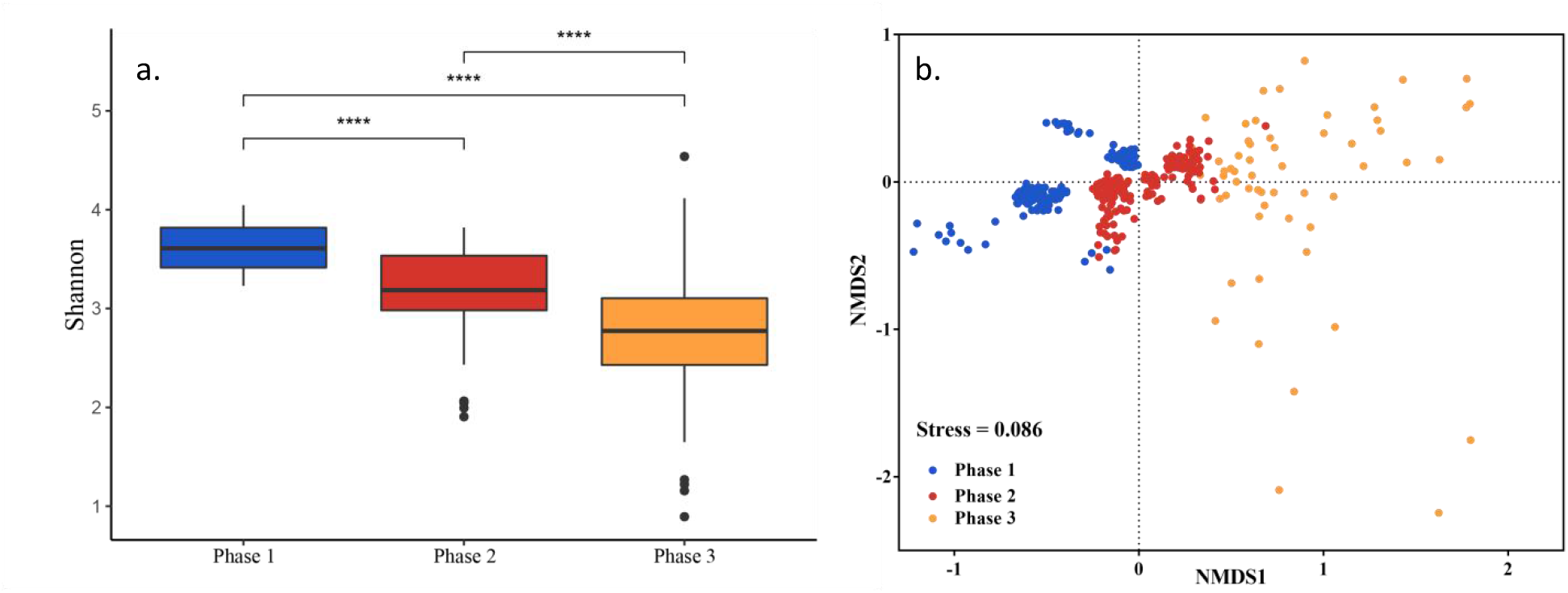
a: Differences in alpha diversity based on the Shannon index (Wilcoxon rank sum test, p< 0.0001). b: NMDS analysis based on Bray–Curtis dissimilarity, showing differentiation in Phases 1-3 (stress = 0.0086).

### Phased changes in the functional traits of microcosm systems

The ASVs of non-acid-producing bacteria at each volume rapidly went extinct. The extinction rate was low in the 1000 ml microcosm system only at 3-5 day (Fig. 5, a). Acid-producing bacterial ASVs did not fluctuate significantly before 35 days but gradually increased in Phase 3 (Fig. 5, b).

**Fig. 5.**
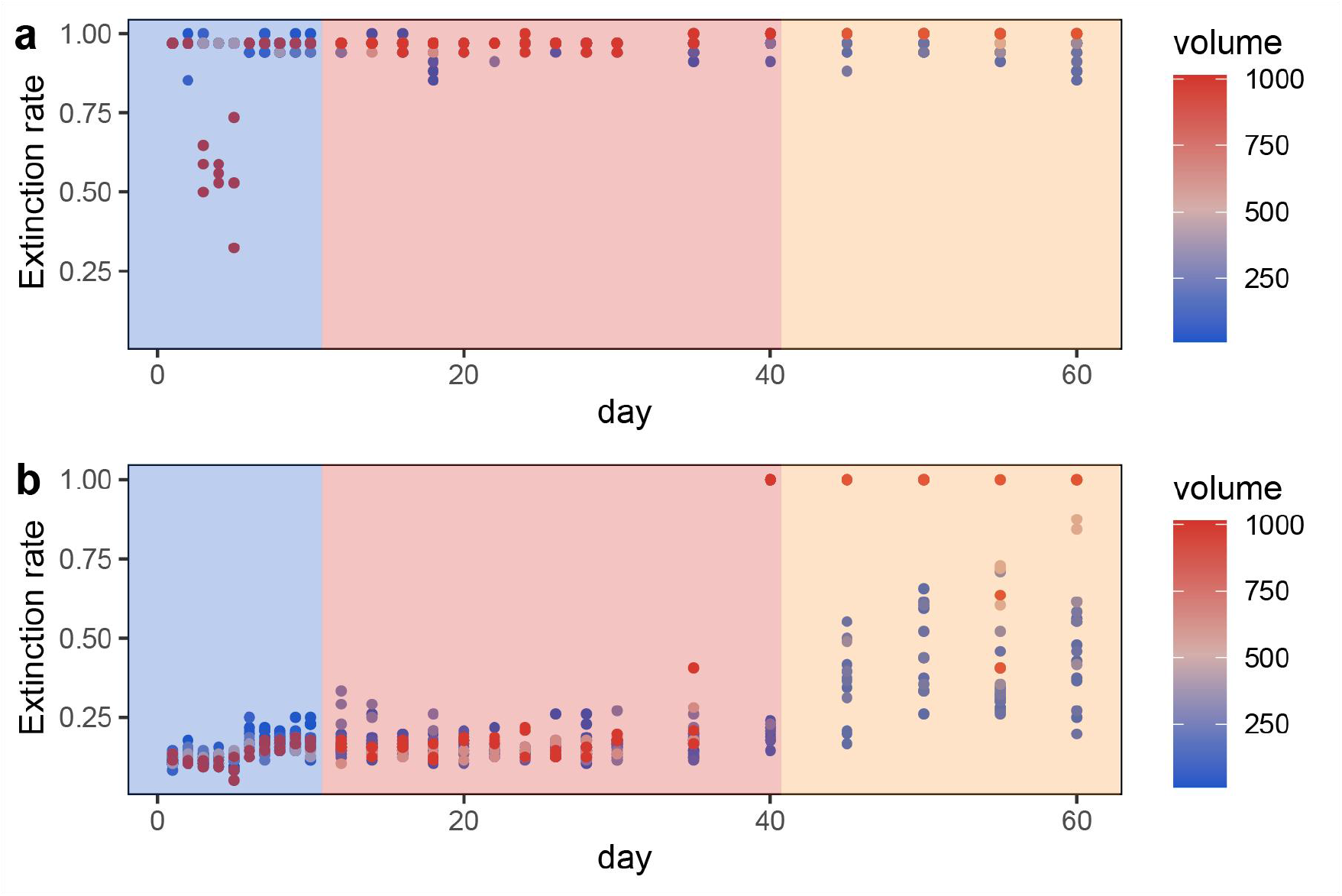
a: Extinction rate of non-acid-producing bacteria. b: Extinction rate of acid-producing bacteria. The colour depth of the dots represents the volume of the microcosm system, increasing in volume from blue to red. The blue background represents Phase 1, red represents Phase 2, and yellow represents Phase 3.

Functional gene prediction revealed two genes related to antibiotic drug resistance, two anti-metabolic bacteriostatic drug resistance genes, and two anti-cancer drug resistance genes. Taking beta lactam resistance, anti-folate resistance, and platinum drug resistance genes as examples, the relative abundance of the three types of genes showed different correlation trends with the volume of the microcosm system at different times. Beta lactam resistance and anti-folate resistance genes had a negative correlation with microcosm volume on 3-5 day of Phase 1 and day 22 and 24 of Phase C, which also showed an SAR (Fig. 6). On 22-30 day, when Phase 2 showed an SAR, the relative abundance of the platinum drug resistance gene tended to be negatively correlated with the volume of the microcosm (Fig. 6).

**Fig. 6.**
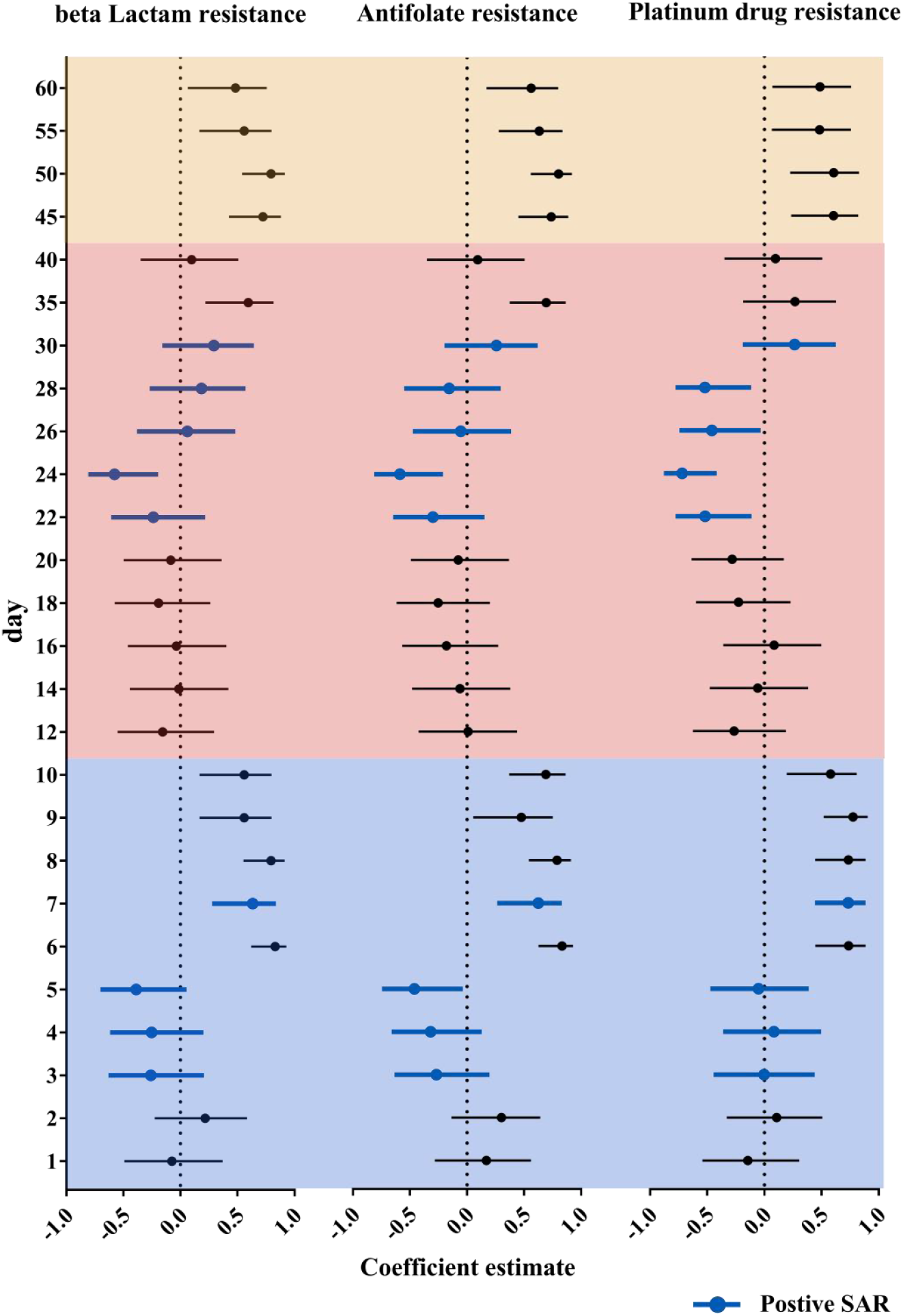
Correlation of the relative abundance of beta lactam resistance, antifolate resistance and platinum drug resistance genes with the volume of the microcosm systems. Correlation coefficients and p values are shown in Table S3. The horizontal axis is the correlation coefficient, and the vertical axis is time. The right side of the dotted line indicates positively correlated, and the left side of the dashed line indicates negatively correlated. Blue corresponds to Phase 1, red corresponds to Phase 2, and yellow corresponds to Phase 3.

## Discussion

This study is the first to find that the occurrence of an SAR is discontinuous over time. The microcosm system had an SAR on days 3-5, 7, and 22-30, accounting for 34.61% of the observation time points and 21.67% of the entire observation time period. The SAR which could be considered as the ecological “law” to some extent, generally ignores the time what is one of the three key elements (time, space and scale) of the ecological research. In recent years, researchers have proposed the species-time-area relationship (STAR) to perfect the SAR, taking into account the temporal dynamics of species richness ^28^. However, the STAR still assumes that an SAR will exist continuously in a time series.

When an SAR can be observed in microcosm systems, the extinction rate and the volume of the system show a negative correlation. This shows that in the closed microcosm, extinction was the main mechanism driving the SAR, which is fully consistent with our hypothesis. Surprisingly, extinction also led to the disappearance of the SAR. The mass extinctions on day 6 and day 40 wiped out previously stable SARs (the extinction rate on day 6 increased by 27.43% from that on the previous day, and the extinction rate on day 40 increased by 12.50% from that on day 35). The timing of these two mass extinctions coincided with the change in pH of the microcosm environment. The process by which biomes change with the environment has a strong consensus in ecology, but there are few examples from ecological studies. This study simply observed the coupling of biomes and their environments. There is no doubt that the decline in environmental pH was driven by acid-producing taxa in the pao cai microbial community ^29^; in turn, the alpha and beta diversity of the microbial communities presented three phases divided by shifts in pH. Arguably, antagonism, or civil war between microbes, led to extinction throughout the experimental period and drove the succession of the microcosm ecosystems. In Phase 1, the acid-producing microbial taxa rapidly lowered the environmental pH, leading to the extinction of the non-acid-producing taxa. The emergence of antibiotic drug resistance genes, anti-metabolic bacteriostatic drug resistance genes and anti-cancer drug resistance genes represents the generation of antagonistic metabolites in the microcosmic system and the potential for extinction of some taxa due to mutual antagonism of microorganisms in the microcosmic system. The extinction caused by antagonistic metabolites was manifested throughout the experimental cycle, and the relative abundance of various antagonistic metabolite suppressor genes appeared to be correlated with the planet volume at the time of SAR production, respectively.. Each full successional stage should include an adjustment stage (early stage), a stabilization stage (middle stage) and a collapse stage (terminal stage). In the early stages of each phase, the gradual occurrence of extinctions regulated the community, and in the middle stage, it stabilized the community. The SAR occurred during the stable stage of the community. As environmental changes continued to accumulate, a mass extinction process occurred at the end of each phase, prompting community succession to the next stage and causing the SAR to disappear until the next stabilization stage. Phase 2 observed in this study should be considered such a complete successional stage.

In general, in our system, extinction shaped SARs but also destroys them; the SARs are discontinuous in time and exist only during stages of ecosystem stabilization. These findings are very important for solving the ecological problems related to SARs and can also be a good reference for sustainable development science. First, previous studies have found that animal and plant SARs are prevalent, while microbial SARs are difficult to find or weak ^30^, and this taxonomic difference is difficult to explain. This study found that an SAR only existed in the stable stage. In microbes, the life cycle is short, community succession is fast, and the response after perturbation is rapid ^31^. Of course, a survey of a particular time point cannot guarantee that a stable stage will be observed, such that no SAR or a very weak SAR would be detected. Animals and plants are more likely to exhibit stable SARs than microorganisms. This is because 1) animal and plant studies usually focus on ecosystems that are less disturbed by humans and are more stable, 2) the longer growth cycle of animals and plants themselves makes their communities stable for a longer period of time, and 3) the rescuer effect brought about by migration and diffusion in open systems offsets the effect of local extinction ^32,33^. Second, the life cycles of microbes are short, and the community succession can be observed over a short amount of time. Thus, in the future, microorganisms should become important taxa for explaining the SAR mechanism and community development. Strengthening research on the temporal dynamics of SARs, starting with species generation, extinction and dispersal, will be critical to clarifying the formation mechanism of SARs. Finally, since SARs only occur during the stabilization stage, the presence or absence of an SAR should serve as an indicator of ecosystem stability. We predict that an SAR will not exist during phases of rapid urbanization, cyanobacterial outbreaks, biological invasion, and other disruptions when biodiversity is plummeting, nor should an SAR exist in the history of major biodiversity explosions and mass extinctions. The gradual extinction of species/taxa is a normal process of ecosystem regulation, but the accumulation of environmental changes in biomes will cause the irreversible collapse of the original ecosystem in the form of mass extinction. This process is similar to the development of human society. Our impact on Earth’s environment is changing the process of extinction ^34,35^. In the future, we should focus on controlling the drivers of extinction within communities in the conservation and management of ecosystems. A complete understanding of its spatiotemporal formation mechanisms will make the SAR a true ecological law.

## Data availability statement

The raw sequence data reported in this paper have been deposited in the Genome Sequence Archive (Genomics, Proteomics & Bioinformatics 2021) in National Genomics Data Center (Nucleic Acids Res 2022), China National Center for Bioinformation / Beijing Institute of Genomics, Chinese Academy of Sciences (GSA: CRA008411) that are publicly accessible at https://ngdc.cncb.ac.cn/gsa/s/L6384eam.

## Conflict of interest statement

The authors declare no competing interests.

## Funding statement

This work was funded by the Second Tibetan Plateau Scientific Expedition and Research Program (STEP), Grant No. 2019QZKK2002.

## Acknowledgements

Thanks to the followings for their contributions in laboratory research, they are Liu li-lei (刘李蕾), Yuan Cai-lian (袁彩莲), Bai Nong-en (白农恩), Yu Guo-bin (俞国宾), Jiang Lin (姜林).

## Notes

### Competing Interest Statement

The authors have declared no competing interest.

