## supplemental information for "Extinction drives a discontinuous temporal pattern of species-area relationships in a microbial microcosm system"

**Supplementary Information**

**Supplementary**
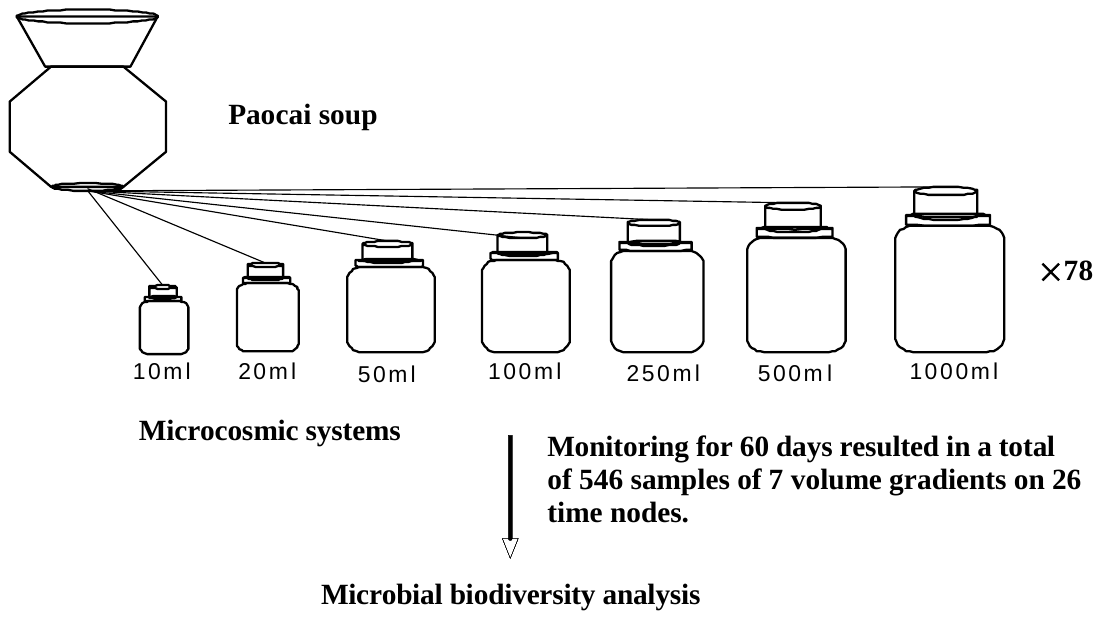
 **Fig. 1 Establishment of the microcosm system.**

Supplementary Table 1 Statistical test results of microbial SAR curves

| day | R2 | p |  | day | R2 | p |
| --- | --- | --- | --- | --- | --- | --- |
| 1 | 0.0466 | 0.3477 |  | 18 | <0.0001 | 0.9810 |
| 2 | 0.0182 | 0.5604 |  | 20 | 0.0087 | 0.6868 |
| 3 | 0.4842 | 0.0005*** |  | 22 | 0.4580 | 0.0008*** |
| 4 | 0.4202 | 0.0015** |  | 24 | 0.2417 | 0.0236* |
| 5 | 0.3273 | 0.0067** |  | 26 | 0.6539 | <0.0001*** |
| 6 | 0.0139 | 0.6109 |  | 28 | 0.6477 | <0.0001*** |
| 7 | 0.3089 | 0.0089** |  | 30 | 0.6031 | <0.0001*** |
| 8 | 0.0817 | 0.2093 |  | 35 | 0.4391 | 0.0011** |
| 9 | 0.0916 | 0.1823 |  | 40 | <0.0001 | 0.9365 |
| 10 | 0.0040 | 0.7849 |  | 45 | 0.163 | 0.0775 |
| 12 | 0.0160 | 0.5847 |  | 50 | 0.1202 | 0.1237 |
| 14 | 0.0033 | 0.8053 |  | 55 | 0.2015 | 0.0412* |
| 16 | 0.0057 | 0.7451 |  | 60 | 0.01358 | 0.6150 |

* p < 0.05, ** p < 0.01, *** p < 0.001

Supplementary Table 2 Statistical test result of the correlation between extinction rate and volume

| day | r | p |  | day | r | p |
| --- | --- | --- | --- | --- | --- | --- |
| 1 | -0.1598 | 0.4889 |  | 18 | -0.1472 | 0.5244 |
| 2 | -0.3393 | 0.1324 |  | 20 | -0.1547 | 0.5032 |
| 3 | -0.6501 | 0.0014** |  | 22 | -0.6844 | 0.0006*** |
| 4 | -0.5659 | 0.0075** |  | 24 | -0.5127 | 0.0175* |
| 5 | -0.5152 | 0.0168* |  | 26 | -0.8193 | <0.0001*** |
| 6 | -0.3464 | 0.1240 |  | 28 | -0.8135 | <0.0001*** |
| 7 | -0.1967 | 0.3928 |  | 30 | -0.7717 | <0.0001*** |
| 8 | 0.03831 | 0.8691 |  | 35 | 0.5124 | 0.0176* |
| 9 | 0.00458 | 0.9843 |  | 40 | -0.0027 | 0.9906 |
| 10 | -0.3874 | 0.0827 |  | 45 | -0.5235 | 0.0149* |
| 12 | 0.1866 | 0.4180 |  | 50 | -0.6306 | 0.0022** |
| 14 | -0.2890 | 0.2039 |  | 55 | 0.4600 | 0.0359* |
| 16 | -0.1860 | 0.4195 |  | 60 | -0.0543 | 0.8151 |

* p < 0.05, ** p < 0.01, *** p < 0.001

Supplementary Table 3 Correlation between the relative abundance of beta Lactam resistance, Antifolate resistance and Platinum drug resistance genes and the volume of microsystems

|  | **Beta Lactam resistance** | |  | **Antifolate resistance** | |  | **Platinum drug resistance** | |
| --- | --- | --- | --- | --- | --- | --- | --- | --- |
| **day** | **r** | **p** |  | **r** | **p** |  | **r** | **p** |
| **1** | **-0.07477** | **0.7474** |  | **0.1705** | **0.4599** |  | **-0.1451** | **0.5302** |
| **2** | **0.2168** | **0.3325** |  | **0.3023** | **0.1716** |  | **0.1075** | **0.6338** |
| **3** | **-0.2592** | **0.2698** |  | **-0.2687** | **0.252** |  | **-0.00182** | **0.9939** |
| **4** | **-0.2523** | **0.2699** |  | **-0.3194** | **0.1581** |  | **0.08336** | **0.7194** |
| **5** | **-0.3872** | **0.0829** |  | **-0.4613** | **0.0353*** |  | **-0.05209** | **0.8226** |
| **6** | **0.8297** | **<0.0001****** |  | **0.8329** | **<0.0001****** |  | **0.7361** | **0.0001***** |
| **7** | **0.6333** | **0.0021**** |  | **0.625** | **0.0025**** |  | **0.7335** | **0.0002***** |
| **8** | **0.7946** | **<0.0001****** |  | **0.7892** | **<0.0001****** |  | **0.7354** | **0.0001***** |
| **9** | **0.5591** | **0.0084**** |  | **0.4755** | **0.0294*** |  | **0.7766** | **<0.0001****** |
| **10** | **0.5591** | **0.0084**** |  | **0.6907** | **0.0005***** |  | **0.5789** | **0.006**** |
| **12** | **-0.1564** | **0.4984** |  | **0.009825** | **0.9663** |  | **-0.2651** | **0.2455** |
| **14** | **-0.01384** | **0.9525** |  | **-0.06209** | **0.7892** |  | **-0.05839** | **0.9015** |
| **16** | **-0.03518** | **0.8797** |  | **-0.1794** | **0.4365** |  | **0.08576** | **0.7117** |
| **18** | **-0.1911** | **0.4067** |  | **-0.2527** | **0.2691** |  | **-0.2254** | **0.3258** |
| **20** | **-0.08462** | **0.7153** |  | **-0.07659** | **0.7414** |  | **-0.2822** | **0.2152** |
| **22** | **-0.2373** | **0.3004** |  | **-0.2984** | **0.1889** |  | **-0.5194** | **0.0158** |
| **24** | **-0.5779** | **0.0061**** |  | **-0.5887** | **0.005**** |  | **-0.7196** | **0.0002***** |
| **26** | **0.06125** | **0.792** |  | **-0.05442** | **0.8148** |  | **-0.4587** | **0.0365*** |
| **28** | **0.1839** | **0.4249** |  | **-0.1557** | **0.5002** |  | **-0.5209** | **0.0155*** |
| **30** | **0.2931** | **0.1972** |  | **0.2573** | **0.2622** |  | **0.264** | **0.2475** |
| **35** | **0.5941** | **0.0045**** |  | **0.6942** | **0.0005***** |  | **0.2674** | **0.2412** |
| **40** | **0.09809** | **0.6723** |  | **0.09461** | **0.6833** |  | **0.09696** | **0.6759** |
| **45** | **0.7239** | **0.0002***** |  | **0.7392** | **0.0001***** |  | **0.6054** | **0.0036**** |
| **50** | **0.7935** | **<0.0001****** |  | **0.8033** | **<0.0001****** |  | **0.6074** | **0.0045**** |
| **55** | **0.5581** | **0.0086**** |  | **0.6335** | **0.002**** |  | **0.483** | **0.0265*** |
| **60** | **0.483** | **0.0266*** |  | **0.5623** | **0.008**** |  | **0.4868** | **0.0252*** |
